# Oligodendrocyte origin and development in the zebrafish visual system

**DOI:** 10.1101/2022.02.14.480410

**Authors:** Adrián Santos-Ledo, Cristina Montes-Perez, Laura DeOliveira-Mello, Rosario Arévalo, Almudena Velasco

**Author notes:** Corresponding author: Almudena Velasco, Instituto de Neurociencias de Castilla y León (INCyL). Universidad de Salamanca., Pintor Fernando Gallego, 1, 37007 Salamanca (SPAIN). These authors contributed equally (Adrián Santos-Ledo, Cristina Montes-Perez).

## Abstract

Oligodendrocytes are the myelinating cells in the central nervous system (CNS). Their developmental origin and specification are well known in birds and mammals but remains unclear in fish. To fulfill this gap, we have studied their early progression during zebrafish visual morphogenesis using the transgenic line Olig2:GFP. We have tracked the Olig2+ cells in the visual system from 48 hours post fertilization (hpf) until 11 days post fertilization (dpf). We have also analyzed the differential expression of the Sox2 and Sox10 transcription factors in this cellular line. Oligodendrocyte progenitor cells (OPCs) originate at 48hpf in regions close to the preoptic area, near to the ventral hypothalamus. Then, at 5 dpf, they migrate to the optic chiasm, where they invade the optic nerve, extending towards the retina. While OPCs in the retina also express Sox2, in the the optic tectum they express Sox10. Within the optic nerve tract, they express both. We have also observed that these Olig2:GFP line do not colocalize with the expression of Mbpa, a myelin marker, but are intimately intertwined. Our data matches with other animal models, where OPCs are specified in the preoptic area and migrate to the optic nerve through the optic chiasm. We revealed that oligodendrocyte is a complex population of cells expressing different transcription factors.

## INTRODUCTION

Oligodendrocytes are the myelinating cells in the Central Nervous System (CNS) of vertebrates (Baumann and Pham-Dinh, 2001; Czopka, 2015; Miller, 2002). They form the myelin sheath, required for the fast saltatory conduction of nerve impulse (Bai et al., 2011; Chung et al., 2013). The visual system, as an intrinsic part of the CNS, is not an exception to myelination (Reichenbach and Bringmann, 2015). Projections within the retina are not myelinated in most mammals, including humans, and its aberrant myelination is associated to age and causes several conditions (Fitzgibbon and Nestorovski, 1997; Reichenbach and Bringmann, 2015). Deficient myelinization specifically in the visual system also lead to several diseases known as Neuromyelitis Optica Spectrum Disorder (Ferilli et al., 2021). In some other vertebrates like teleosts, birds or reptiles, the optic nerve (ON) inside the retina is partially or fully myelinated (Lillo et al., 1998; Münzel et al., 2012; Parrilla et al., 2016). Despite these differences, oligodendrocytes share many key transcriptional factors and specification routes between mammals and fish (Buckley et al., 2008; Lyons and Talbot, 2015; Mathews and Appel, 2016; Preston and Macklin, 2015).

Oligodendrocytes arise from the Oligodendrocyte Progenitor cells (OPCs), which are characterized by the expression of several transcription factors, including Olig2, Sox2 and Sox10 (Mathews and Appel, 2016; Park et al., 2002; Takada et al., 2010). OPCs are highly mobile and proliferative cells, which migrate and search for their target axons along the nervous tracts until they ensheathe them. In that moment they become myelinating oligodendrocytes and start expressing, along with Olig2 and Sox10, myelin proteins such as Mbpa, Mpz, or Claudin κ (Bai et al., 2011; Brösamle and Halpern, 2002; Jung et al., 2010; Mathews and Appel, 2016; Münzel et al., 2012; Nawaz et al., 2013). In vertebrates like chickens and mice, OPCs in the visual system are originated in the preoptic area (Gao and Miller, 2006; Merchán et al., 2007; Miller, 2002; Ono et al., 2017), from where they migrate to the optic chiasm (Bribián et al., 2006; Gao and Miller, 2006; Merchán et al., 2007; Ono et al., 1997). However, the place where this OPCs originate in teleosts is not clear (Tian et al., 2016). Furthermore, the myelinization process during development is not fully understood. For example, although the first evidence of myelinization (myelin binding protein a, *mbpa*, expression) are reported from 2 dpf, the entrance of mature oligodendrocytes in the retina does not occur until much later (Brösamle and Halpern, 2002; Buckley et al., 2010).

Both Olig2 and Sox10 are expressed in other regions outside the CNS and more interestingly OPCs express other members from the Sox family. For example, Sox2 has implicated in the development of the visual system (Graham et al., 2003; Reinhardt et al., 2015). Sox2 controls proliferation and cell fate (Mercurio et al., 2019). Strikingly, while Sox10 remains in mature oligodendrocyte, Sox2 only remains in some oligodendrocytes and neurons. Its function in differentiated cells is still a matter of debate (DeOliveira-Mello et al., 2019).

Our work intends to clarify the developmental origin and specification of OPCs in the zebrafish visual system, defining the characteristics of these cells, the expression pattern in relation to Sox proteins, as well as the onset of myelination of visual tracts. We show the origin of OPCs in the preoptic area at 2dpf, and how they penetrate the optic nerve from the optic chiasm. We also stablished the onset of myelination at 5 dpf in the optic chiasm, which spread dorsal- and ventrally to the rest of the visual system. We also show that some cells retain Olig2 and Sox2 expression within the retina but the oligodendrocytes in the brain present Sox10 instead. The OPCs that invade the optic nerve present both transcription factors. Finally, to analyze early myelinization we used the Mbpa:RFP transgenic line and we found a different population that express this marker but not Olig2. This reveals the complex identity of the oligodendrocytes.

## MATERIAL AND METHODS

### Animals

Zebrafish embryos were obtained by natural mattings. Eggs were raised in E3 medium at 28.5ºC, and collected at different stages, according to *Kimmel et al., 1995*. We employed several transgenic lines: *Tg(olig2:EGFP)* (Shin et al., 2003), *Tg(sox10:GFP)* (Dutton et al., 2008), *Tg(sox10:tagRFPt)* (Blasky et al., 2014), *Tg(mbpa:tagRFPt)* (Ravanelli et al., 2018) and *Tg(gfap:EGFP)* (Bernardos and Raymond, 2006). All specimens were deeply anesthetized in tricaine methanesulfonate before sacrifice, according to Spanish and European laws (2010/63/EU; RD 53/2013; Ley 32/2007; Orden ECC/566/2015).

All protocols were performed according to the European Union Directive 86/609/EEC and Recommendation 2007/526/EC, regarding the protection of animals used for experimental and other scientific purposes, enforced in Spanish legislation under the law 6/2013. All protocols were approved by the Bioethics Committee of the University of Salamanca

For each stage and staining at least 10 embryos from three different parents were used.

### Embryo manipulation

For tissue sections, Embryos/larvae were hand-dechorionated when necessary and fixed in 4%paraformaldehyde in 0.1M pH 7.4 phosphate buffer saline (PBS) overnight at 4ºC. After three washes in PBS, embryos were embedded in a solution containing 10% sucrose and 1.5% agarose. Then blocks were cryoprotected in 30% sucrose in PBS overnight at 4ºC. 12μm coronal sections were obtained in a cryostat (Thermo Scientific HM560).

For life imaging, embryos at the proper stage were anesthetized as usual and embedded in 1.5% low melting agarose (ThermoFisher Scientific R0801) in E3 medium.

### Immunohistochemistry

Sections were washed several times in PBS and then incubated for 90 minutes in 5% normal donkey (DK) serum in PBS with 0.2% Triton X-100 at room temperature (RT). After that, primary antibodies (Table 1) were incubated overnight in 5% normal DK serum, 0.2% Triton X-100 and 1% dimethyl sulfoxide (DMSO) at 4°C. Sections were washed in PBS and then incubated 90 minutes at RT in darkness with a 1:400 dilution of Alexa 488, Alexa 555 or Alexa 647 fluorescent secondary antibodies (Table 2), in a buffer containing 5% normal DK serum in PBS. Next, sections were washed in PBS and then incubated for 10 minutes in 1:10000 4’,6-diamidino-2-fenilindol (DAPI; Sigma) for nuclei counterstaining. Sections were washed thoroughly and mounted with Fluoromount-G® Mounting Medium (Invitrogen).

**Table 1.**
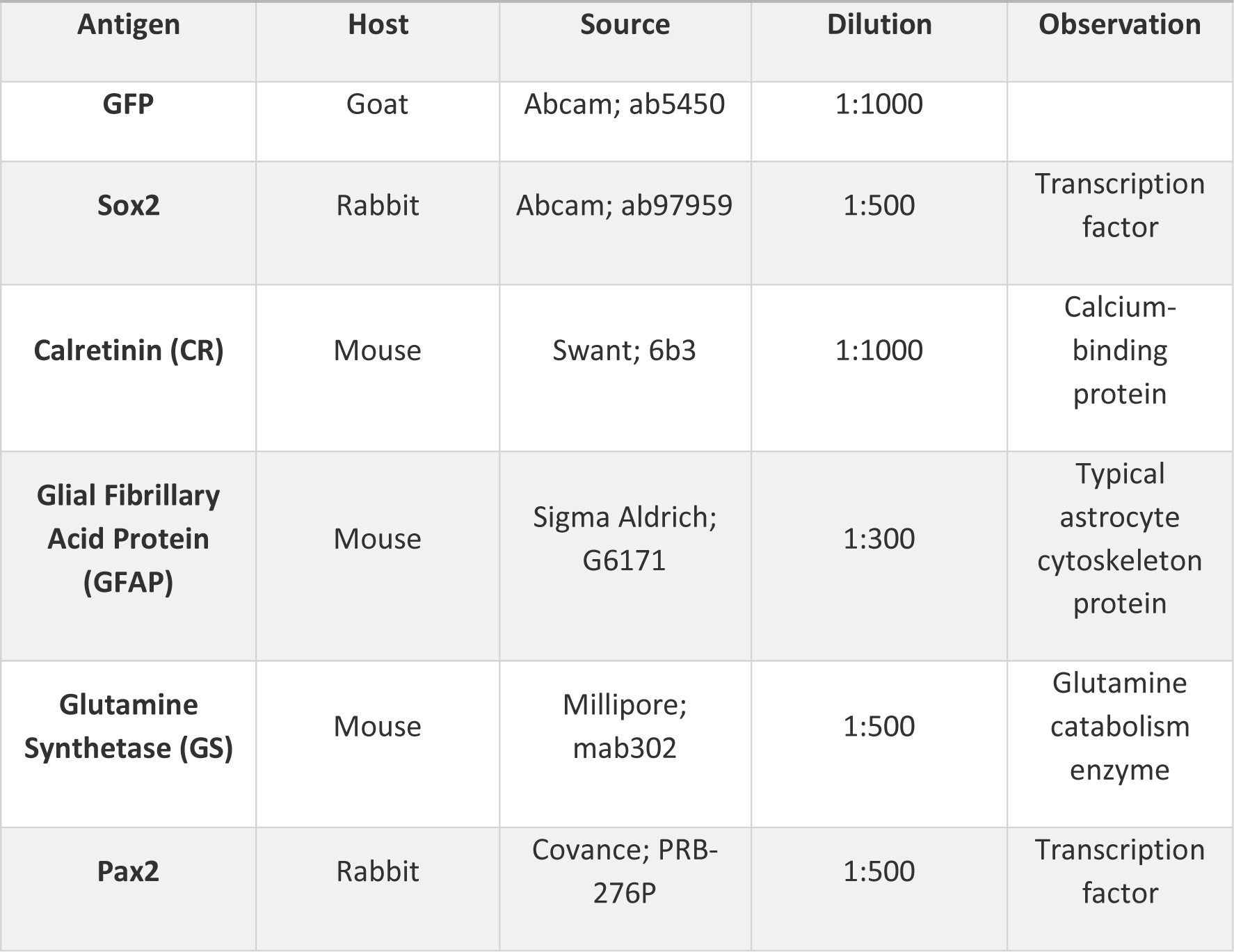
Primary Antibodies.

| Table 1. Primary Antibodies |  |  |  |  |
| --- | --- | --- | --- | --- |
| Antigen | Host | Source | Dilution | Observation |
| GFP | Goat | Abcam; ab5450 | 1:1000 |  |
| Sox2 | Rabbit | Abcam; ab97959 | 1:500 | Transcription factor |
| Calretinin (CR) | Mouse | Swant; 6b3 | 1:1000 | Calcium-binding protein |
| Glial Fibrillary Acid Protein (GFAP) | Mouse | Sigma Aldrich; G6171 | 1:300 | Typical astrocyte cytoskeleton protein |
| Glutamine Synthetase (GS) | Mouse | Millipore; mab302 | 1:500 | Glutamine catabolism enzyme |
| Pax2 | Rabbit | Covance; PRB-276P | 1:500 | Transcription factor |

**Table 2.**
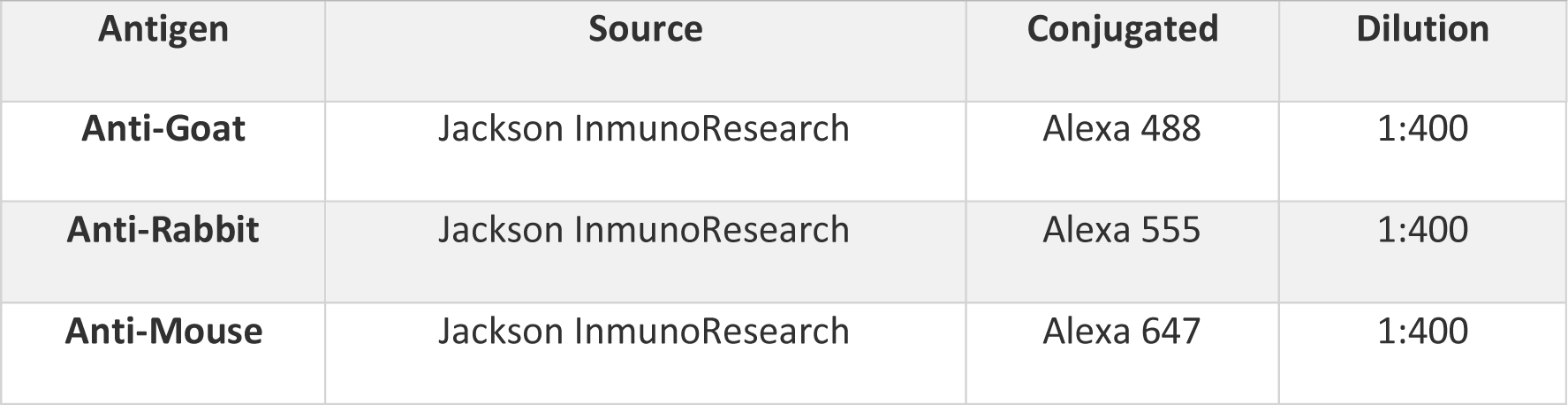
Secondary Antibodies.

### Image acquisition

Images were obtained with a *Leica Stellaris* (inverted DMI8) microscope. Living embryos (Mbpa:RFP/Olig2:GFP double transgenic line) were imaged with *Leica Stellaris* (inverted DMI8) microscope using a 20x objective from the dorsal side. Sections were imaged using a 40x oil immersion objective. In case of the sections four tiles were acquired and automatically assembled by the software. Acquired z-stacks were transformed into maximum intensity projections using the Las X software from Leica. Images were later cropped or rotated in Image J. Bright and contrast were only adjusted for better visualization. Finally, figures were built using *Photoshop CS5*.

### Quantification and statistics

Colocalization of different markers was quantified using appropriate plugins in ImageJ. Six embryos were used per stage and the total number of Mbp+ cells in the optic nerve chiasm were counted. ANOVA test with a Bonferroni post-test Statistics were performed using Graphpad Prism

## RESULTS

### OPCs markers are detected from 48 hpf onwards

Oligodendroctye progenitor cells (OPCs) were identified using the Olig2:GFP transgenic line. Although Olig2 was expressed earlier, Sox2 expression was triggered at 48 hpf. In the retina Sox2+ cells were clustered in the peripheral retina and more dispersedly in the central part (Fig 1A, B). Olig2 displayed the opposite pattern (Fig 1A, B) but colocalization between both markers was detected in the external central retina, the prospective external nuclear layer (Fig 1B). We also observed some single Sox2+ cells outlying the optic nerve head (ONH) (Fig 1B). Olig2 was present in the hypothalamus and in the optic tectum but separated from Sox2 (Fig. 1A, C). Sox10 was analyzed using the line Sox10:RFP and it was absent in the retina except from some cells in the optic nerve area (Fig 1D, E). Olig2+ cells in the hypothalamus were positive for Sox10, too (Fig 1D, F). As expected, we also observed single labeled Sox10+ cells in the optic tectum since this transcription factor is expressed in cells other than oligodendrocytes (Fig. 1F).

**Figure 1.**
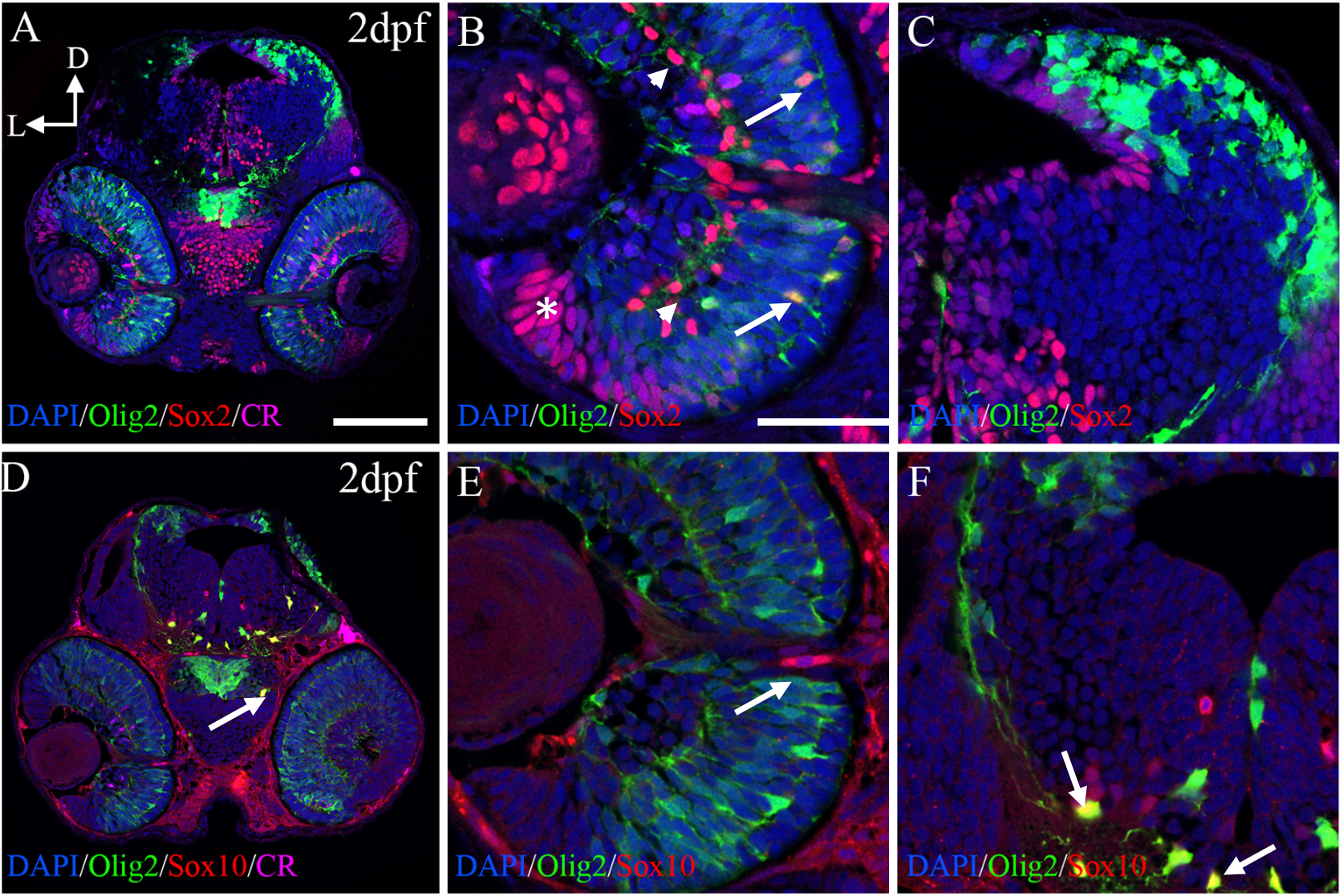
OPCs markers are detected from 2 dpf onwards. Distribution of Olig2+/Sox2+ (A-C) and Olig2+/Sox10+ (D-F). Sox2 and Olig2 colocalize in the central outer retina (arrows in B) but not in the inner (arrowheads in B) or peripheral retina (asterisk in B). Separation between Sox2 and Olig2 also exists in the rest of the brain (C). Sox10 is absent from the retina except cells in the optic nerve (D, E). In the hypothalamus and optic tectum, Olig2+ cells also present Sox10 (arrows in D and F). Calretinin (CR) is used as a marker for visual system differentiation. Scale bar in A, D: 100μm; in B, C, E, F: 50μm

### From 3dpf onwards Olig2+ cells spread throughout the visual system

At 3dpf when retinal lamination is evident in zebrafish, Olig2+/Sox2+ cells were observed in the inner nuclear layer (INL) and in the ganglionar cell layer (GCL) (Fig 2A, B). Olig2+/Sox2+ cells were also detected in ONH (Fig 2B) and the optic nerve chiasm labeled with Calretinin (CR) (Fig 2C). Within the visual system (retina, optic nerve and optic tectum) all Olig2+ cells also presented Sox2. Olig2+/Sox2+ cells were very abundant in the ventricular zones indicating maturing but not fully differentiated cells. In other parts of the brain, such as hypothalamus, Olig2+ cells were negative for Sox2 (Fig 2D). At this stage Sox10+ cells are absent in the retina (Fig 2E, F) but we found a subpopulation of them in the optic nerve chiasm negative for Olig2 (Fig 2G). Olig2+ cells far from the ventricular zones with full arborization indicating mature oligodendrocytes were also positive for Sox10 (Fig 2H). Neither Olig2+/Sox2+ cells nor Sox10+ cells presented ramifications around the optic nerve, indicating that myelinization had not started yet.

**Figure 2.**
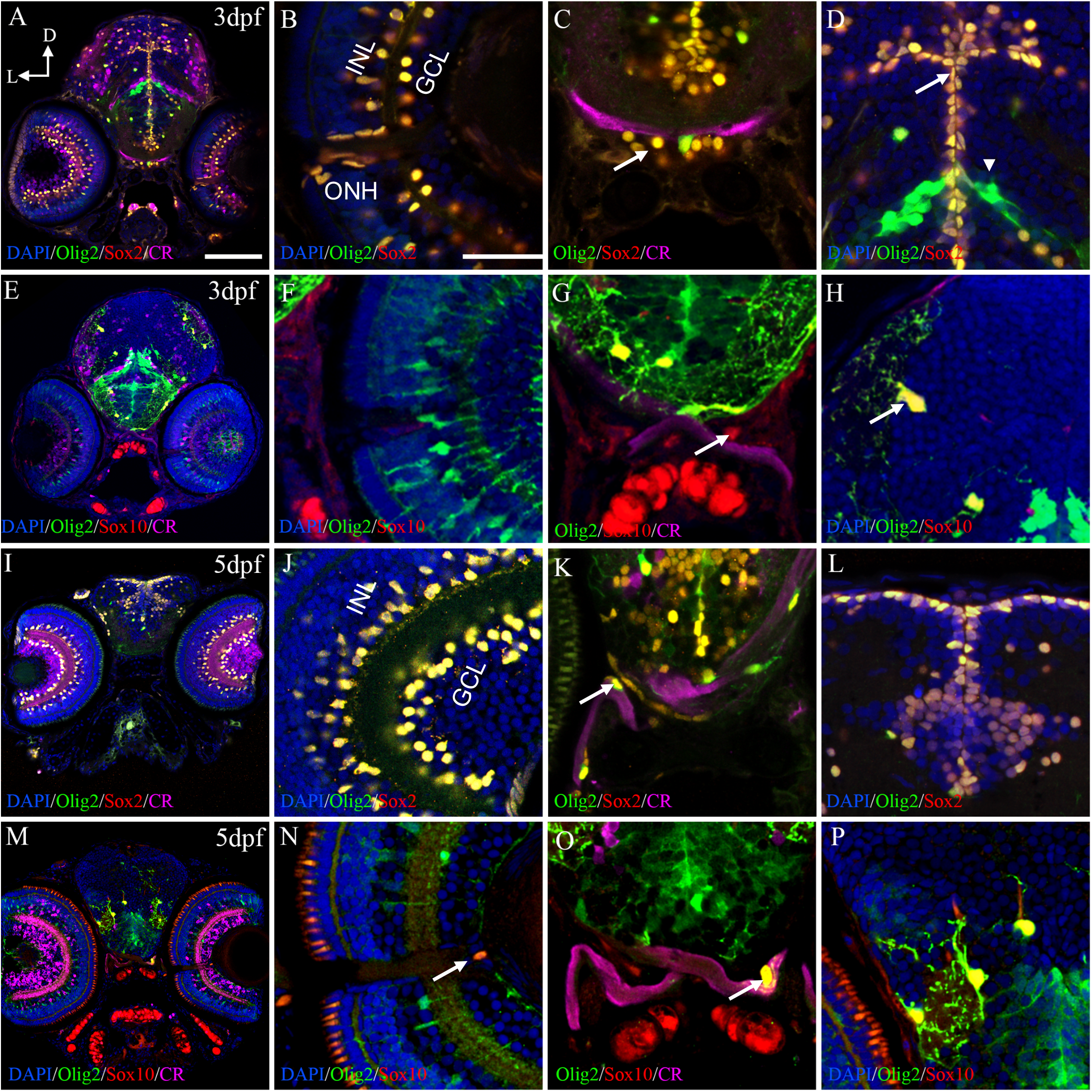
From 3 dpf onwards Olig2+ cells spread throughout the visual system. At 3 dpf, Olig2+/Sox2+ cells are present in the INL, GCL, ONH (A, B), the optic chiasm (arrow in C) and the ventricular zones of the brain (arrow in D). Arborized Olig2+ cells are negative for Sox2 (arrowhead in D). Sox10+ cells are absent in the retina but are present in the optic chiasm (E, F, arrow in G). Fully arborized Oig2+ cells are positive for Sox10 (arrow in H). Sox2 distribution is identical at 5 dpf (I-L). Olig2+/Sox10+ cells are found in the ONH and the optic nerve (M-O) with projections surrounding it (arrow in O.) Fully arborized Olig2+ cells also present Sox10 (P). Calretinin (CR) is used as a marker for visual system differentiation and to label the optic nerve. GCL: ganglionar cell layer; INL: inner nuclear layer; ONH: optic nerve head. Scale bar in A, E, I, M: 100μm; in B, C, D, F, G, H, J, K, L, N, O, P: 50μm

From 5 dpf onwards we detected little to no changes in the distribution of the Olig2+/Sox2+ cells (Fig 2I-L). Olig2+ cells in the retina, optic nerve and optic tectum always express Sox2. However, there were some Olig2+/Sox10+ cells in the ONH (Fig 2M, N) and in the optic nerve chiasm. Furthermore, these latter cells seemed to surround the optic nerve with their projections (Fig. 2O). In the other parts of the brain Olig2+ cells continued expressing Sox10 (Fig 2P).

Since we found Olig2+/Sox2+ cells in several areas of the retina, we wondered if they presented other typical markers for retinal cells. Olig2+ cells also expressed GFAP (Fig 3A-C) and Glutamine synthetase (GS) (Fig 3D-F) indicating that they could be Müller and amacrine cells partially differentiated. We were curious about the Olig2+/Sox2+ in the ONH. The formation of the optic nerve and its exit from the retina towards the optic tectum is still a process not very well understood. Pax2 is very important during this process, but we did not find any Olig2+/Pax2+ cells suggesting that the ONH presents a complex population of cells during early development (Fig 3G-I).

**Figure 3.**
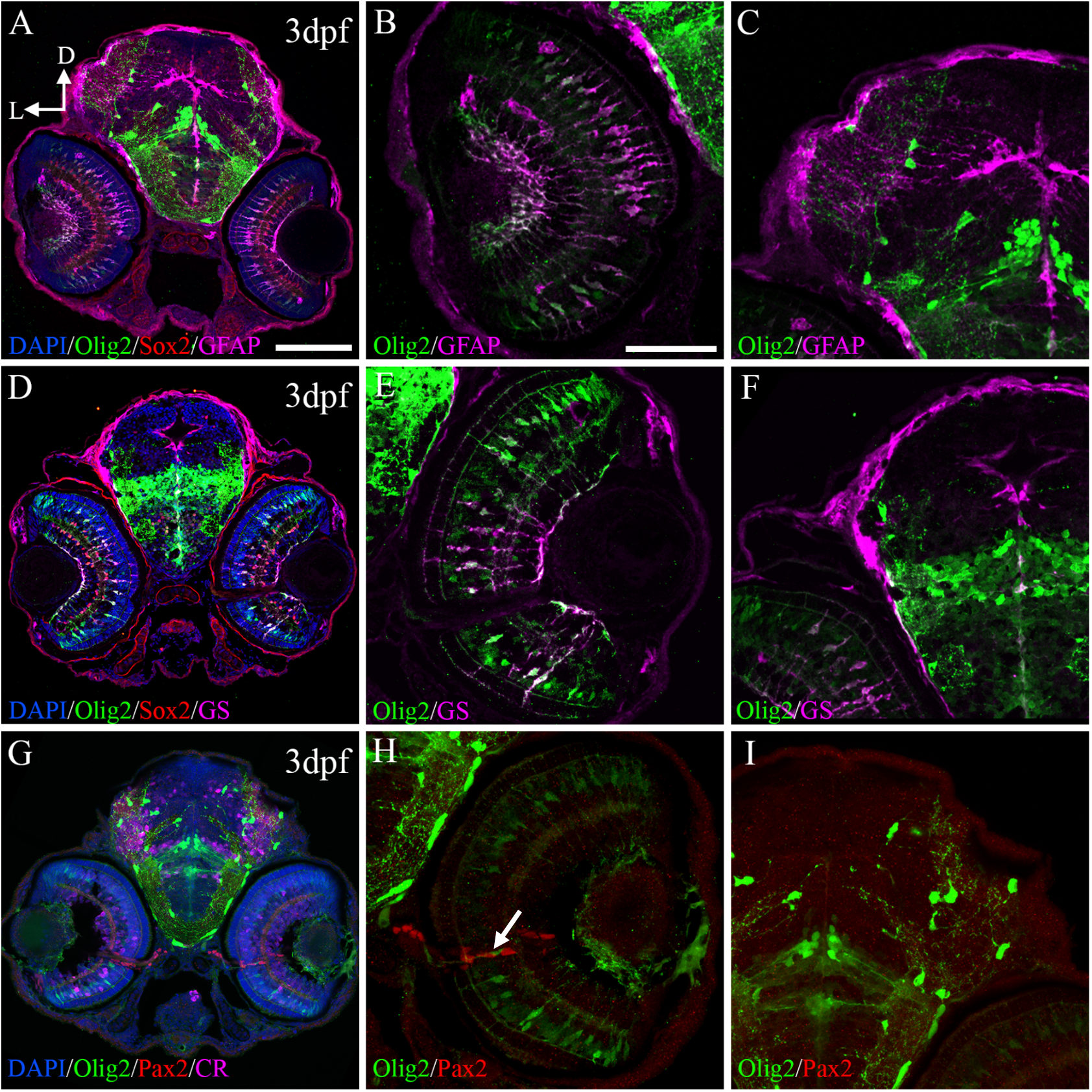
Olig2+ express other retinal markers. Olig2+ cells also express GFAP (A, B) and GS (D, E) in the retina but not in the optic tectum (C, F). Olig2+ cells do not present Pax2 in any area (G-I) but these two populations are intertwined in the ONH (arrow in H). ONH: optic nerve head. Scale bar in A, D, G: 100μm; in B, C, E, F, H, I: 50μm

### Olig2+ cells are fully differentiated from 7 dpf

The changes we started to see at 5 dpf culminated at 7 dpf. In the retina, the optic chiasm, optic tectum and the ventricular areas of the brain, Sox2+ cells were not positive for Olig2 (Fig 4A-D). This would suggest the existence of an important differentiating window for oligodendrocytes between 5 and 7 dpf. In the retina, Olig2+ cells were restricted to the proliferative growth zone (PGZ), where undifferentiated cells exist (Fig 4A, E). Colocalization between Olig2 and Sox10 increased at this stage (Fig 4E). Although they were still not very abundant, Olig2+/Sox10+ were observed in the optic nerve chiasm with projections surrounding the optic nerve (Fig 4G). Arborized Olig2+ cells in the optic tectum retained Sox10 expression (Fig 4H).

**Figure 4.**
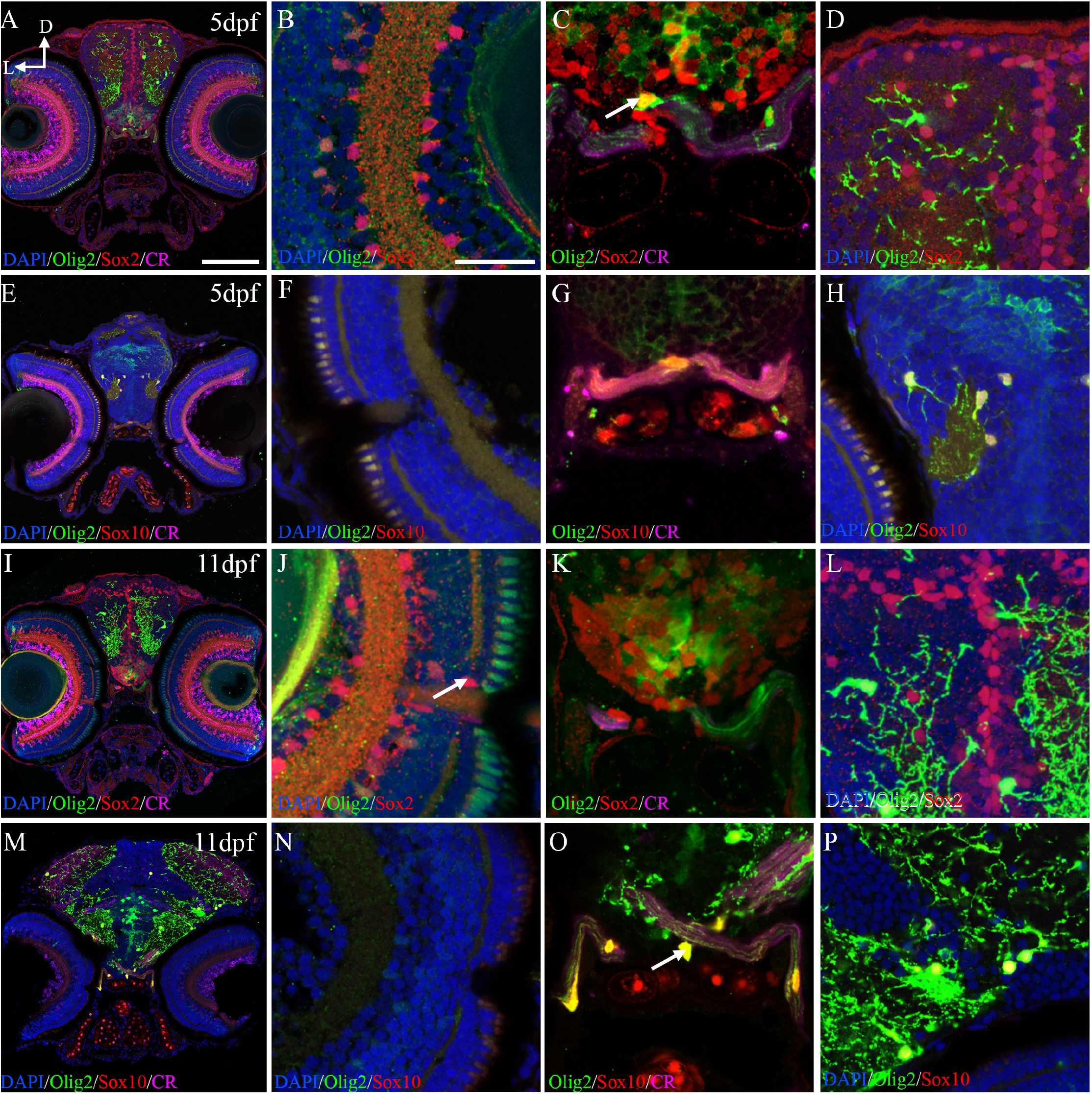
Olig2+ cells are fully differentiated from 7 dpf. At 7 (A-D) and 11 (I-L) dpf, Sox2+ cells generally do not present Olig2. Sox2+ cells are abundant in the retina (A, B, I, J), the ONH (arrow in J) and the proliferative areas of the brain (L). Sox2 occasionally present Olig2 in the optic nerve chiasm at 7 dpf but not at 11 dpf (arrow in C). In the optic tectum, Sox2+ cells have a close relation with Olig2+ cells but they never only colocalize (D, L). Olig2+ cells express Sox10 in the optic nerve chiasm (E, G) and in other parts of the brain (H). Differentiated part of the retina was empty of Olig2+ and Sox10+ cells (F). At 11 dpf, Sox2 is present in the retina and the ONH (I, J) and in the optic nerve although they are not positive for Olig2 (K). In the optic tectum, Sox2+ cells are restricted to the ventricular area (L). Olig2+/Sox10+ cells were abundan in the optic nerve with obvious projections (M, O) and there are none in the retina (N). In the optic tectum Olig2+ cells continue expressing Sox10 (P). ONH: optic nerve head. Scale bar in A, E, I, M: 100μm; in B, C, D, F, G, H, J, K, L, N, O, P: 50μm

Respect to Sox2, no changes were observed at 11 dpf. Retina and ventricular areas presented many Sox2+ cells with no Olig2+ expression (Fig 4A-L). Although we found several Sox2+ in the optic nerve, they were negative for Olig2 (Fig 4K). Olig2+ cells and Sox2+ however presented an intimate relationship (Fig 4L). At 11 dpf, the evidence of colocalization between Olig2 and Sox10 were more evident (Fig. 4M). While the retina did not present positive cells (Fig. 4N), the optic nerve had Olig2+/Sox10+ cells with their projections enseathing the optic nerve (Fig. 4O). In the optic tectum the oligodendrocytes continued expressing Sox10 (Fig. 4P).

### Myelinization of the visual system starts at 5 dpf

One of the most important functions of oligodendrocytes is the myelinization of axons within the central nervous system. To follow up the relation between Olig2 transgene and the myelinization process, we established a new combination with other transgenic line for myelin binding protein a (Mbpa): Olig2:GFP/Mbpa:RFP. The presence of Mbpa in the spinal cord was detected at 4 dpf but in the visual system areas was at 5 dpf (Fig 5A, A’). Even though, only a few Mbpa+ cells were detected in the optic tectum and in the optic nerve. In fact, this latter only presented 2 or 3 Mbpa+ cells (quantified in Fig 5D). We observed a significant increase of Mbpa cells in the optic nerve area from 5 dpf until 11 dpf (Fig. 5A’, B’, C’, quantified in D). Interestingly, we did not detect a clear colocalization between Olig2 and Mbpa in any of the areas. However, we found that Olig2 and Mbpa cells were always in close relation both in the optic tectum (Fig 5A-C and insets) and the optic nerve (Fig 5A’-C’ and insets), partially overlapping. At least 40% of Olig2+ cells in the optic nerve had an associated Mbpa+ cell at 5 dpf, a number that increased significantly up to 60% at 7 dpf (quantified in Fig 5E). This reveals that the Olig2:GFP might not be labeling the first myelinating cells, at least in the visual system.

**Figure 5.**
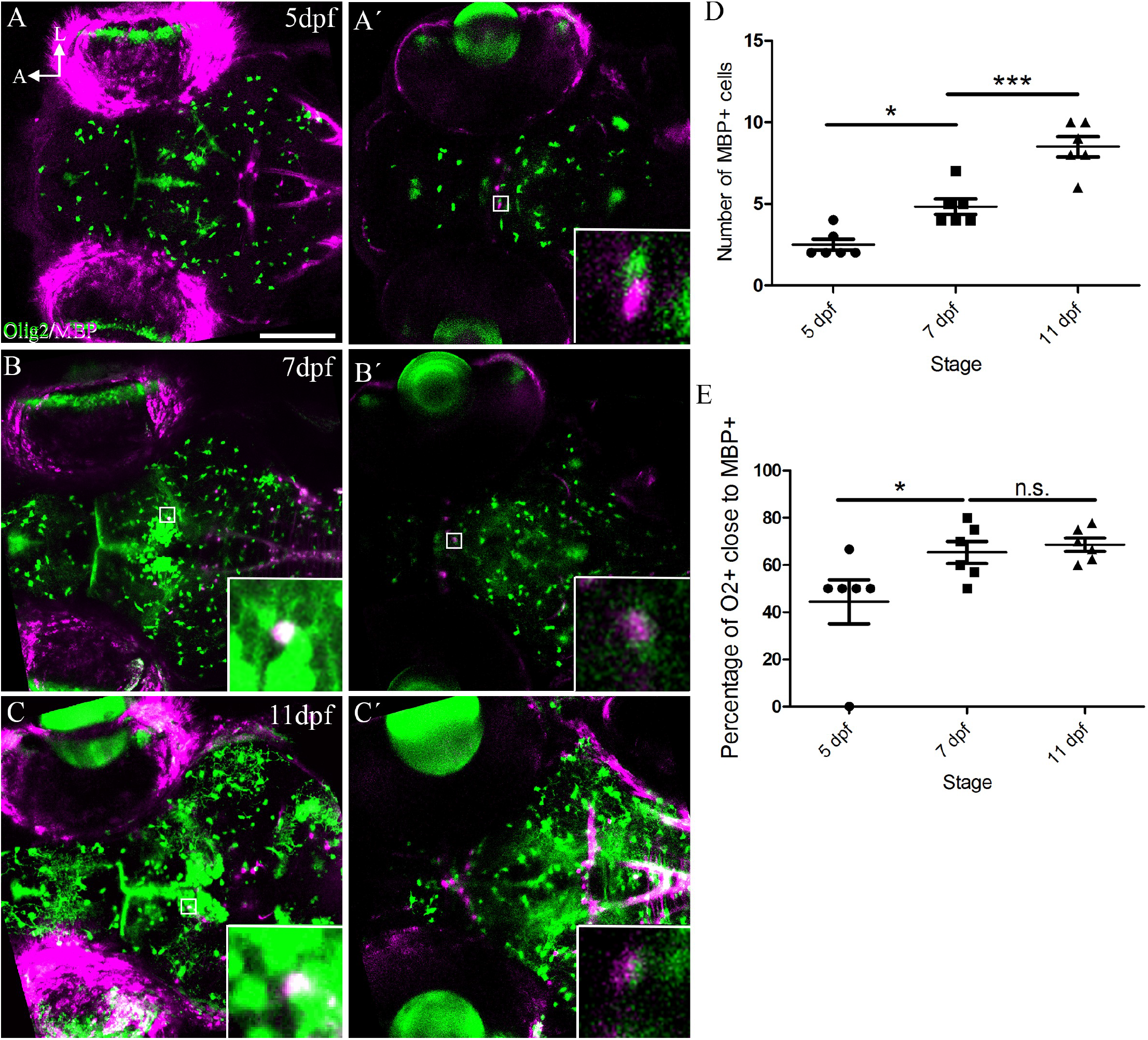
Myelinization of the visual system. Confocal images of life Olig2:GFP/Mbpa:RFP embryos at 5 (A, A’), 7 (B, B’) and 11 (C, C’) dpf. A 25 μm thick stack is shown as max intensity projection in the optic tectum area (A, B, C) and the optic nerve chiasm (A’, B’, C’). Expression of Mbpa starts at 5 dpf in the visual system (A, A’). The number of Mbpa+ cells increase significantly from 5 dpf to 11 dpf (A’, B’, C’, quantified in D). Although we do not observe a strong colocalization, Olig2+ and Mbpa+ cells and intimately associated (insets). The number of associated cells increases from 5 dpf to 7 dpf (quantified in E). Scale bar: 100μm

## DISCUSSION

While the onset of myelination in the spinal cord has been studied in depth (Hines et al., 2015; Kucenas et al., 2009; Mathews et al., 2014; Ravanelli and Appel, 2015; Takada et al., 2010), there is not much known regarding the myelinization in the zebrafish visual system. Tian et al. (2016) analyze this process but starting at 7 dpf and, thus, do not clarify the oligodendrocyte’s origin. Other studies that use ablation or demyelination experiments also describe the formation of oligodendrocytes and their evolution, but they do not pinpoint their origin (Chung et al., 2013; Fang et al., 2015; Münzel et al., 2014).

In zebrafish, we lack a specific marker of oligodendrocytes but the Olig2:GFP and other transgenic lines provide strong and useful information. These lines also have some limitations since they might not be labelling oligodendrocytes at all stages and cells might retain GFP after the expression of the promotor is stopped. Here we show a complex and diverse oligodendrocyte population that change expression and location over time. Oligodendrocyte Progenitor cells (OPCs) specification starts in the preoptic area, near the ventral hypothalamus at 2 dpf. This coincides with data from mice and chicken (Gao and Miller, 2006; Merchán et al., 2007; Miller, 2002; Ono et al., 2017, 1997). These OPCs migrate through the brain until they reach the optic chiasm, from where they finally colonize the optic nerve (Garcion et al., 2001). Our results validate this model also in zebrafish. Furthermore, we have established that this process starts as early as 2 dpf.

We have also found Olig2 in the retina. At early stages (2 to 5 dpf) together with Sox2 and later restricted to the proliferative growth zone (PGZ) where retinal progenitors are. Sox2 is a transcription factor related to neural progenitors (Bylund et al., 2003). At 3dpf, Olig2+ cells also present markers for Müller glia (GS+). This expression has been associated to developmental and regenerative processes, since Müller cells act as retinal neural progenitors (Fimbel et al., 2007; Nakamura et al., 2006; Shibasaki et al., 2007; Thummel et al., 2008). Linked to the onset of myelinization, Olig2+ cells stop expressing Sox2 and trigger Sox10. While oligodendrocytes in the optic tectum always expressed Sox10, this transition is obvious in the optic nerve. This might be a consequence of the transition from the proliferative state of OPCs to more mature oligodendrocytes (Ono et al., 2001). This view is reinforced by our description of Olig2/Sox2 cells in the midline, a typical neurogenic zone (Germanà et al., 2010).

Curiously, we detected some Sox10+ in the optic nerve head (ONH) area at different timepoints. Sox10 is a widely expressed transcription factor from cartilage (Santos-Ledo, et al., 2017) to pericytes involved in the development of blood vessels (Trost et al., 2013). However, zebrafish retina does not contain intraretinal vessels. Sox10 belongs to a big family of transcription factors. In other vertebrates, Sox9 is expressed in the retina (Fischer et al., 2010; Plavicki et al., 2014; Yokoi et al., 2009) and control glial specification (Haldin and LaBonne, 2010; Kordes et al., 2005; Stolt and Wegner, 2010). In zebrafish Sox10 might fulfill this role. Thus, Sox10 may play an unknown role during retinal specification in zebrafish retina. Also, surrounding the ONH, we have found Olig2+/Sox2+ that lose the Olig2 expression over time and, based on our previous experience (Parrilla et al., 2012, 2013), we hypothesized that they could be the reticular astrocytes that form the optic stalk, however, they do not express Pax2 as it happens in mouse and adult zebrafish (Parrilla et al., 2013; Tiwari et al., 2014). One provoking idea is that this Sox2+ cells are progenitors of the reticular astrocytes. Thus, we have found a new type of cells that might contribute to the formation of the optic stalk in zebrafish that require further study.

Myelination, marked by Mbpa+ cells, is first observed at 5dpf in the optic chiasm and in the beginning of the optic nerve. In zebrafish CNS, the Mauthner axons are the first ones to become myelinated by 3dpf (Buckley et al., 2010). *mbpa* is clearly expressed in some brain regions by 4dpf (Brösamle and Halpern, 2002; Jung et al., 2010; Pinzon-Olejua et al., 2017), as well as *plp/DM20* in the optic nerve (Brösamle and Halpern, 2002; Yoshida and Macklin, 2005). However, the same authors did not find myelinated axons in the optic nerve by electron microscopy until 7dpf. In a similar way, another group found mpz axons in the optic nerve at 7dpf (Bai et al., 2011), although *mpz* expression started at 5dpf in the optic chiasm (Schweitzer et al., 2003). All together this data indicate that myelination is a progressive process that might be trigger in different places simultaneously. Our results match with these results, Mbpa+ cells are detected at 5 dpf in the visual system increasing significantly until 11 dpf. Interestingly, we failed to find clear colocalization between Mbpa+ and Olig2+ cells. However, we have found a close relationship between these cells. Thus, during early zebrafish development the first cells expressing Mbpa might not be oligodendrocytes or, as previously suggested by *Hughes and Appel, 2020*, we are detecting processes associated with myelin rather than oligodendrocytes bodies. Unfortunately, we failed to detect the RFP in frozen sections since fluorescence is quench by fixation. Further experiments are needed to characterize this interaction.

At later stages, 7 dpf onwards, we have detected some Sox2+ cells that do not express Olig2 in the optic nerve area. These cells might be specific to zebrafish since its visual system continuously grows, adding new axons to the visual pathway (Johns and Easter Jr., 1977). The remaining Sox2 population might generate new myelinating oligodendrocytes. An idea reinforced by the fact that Sox2+ cells are associated to Olig2+ cells. Axonal selection and myelination by oligodendrocytes are limited to a 5h time lapse (Czopka et al., 2013), despite myelination remains until adulthood (Jung et al., 2010). According to these data, newly formed axons would be myelinated progressively by the OPCs, which could be proliferating as they detect these axons. The other possibility is that the mature oligodendrocytes could myelinate the new axons, leaving the OPCs (Sox2+ cells) as resident repairing cells. However, more studies are needed to clarify this point. Zebrafish, an animal where multiple de- and remyelination experiments events occur might be an ideal model to do so (Chung et al., 2013; Fang et al., 2015; Münzel et al., 2014).

Our results show that OPCs in the zebrafish visual system are generated in similar brain areas to other groups of vertebrates. We have also found that myelination starts at 5dpf in the optic chiasm, and that myelination and oligodendrocytes extend both dorsally and ventrally in the visual pathway. Our results show the progression of OPCs through the optic nerve and optic tracts. Therefore, in zebrafish the visual system myelination in zebrafish resemble that described other vertebrates despite the morphological differences and validate this model to study human diseases related to aberrant myelinization.

## Acknowledgements

We would like to thank Prof. Bruce Appel for the transgenic lines used in this work. We also would like to thank Consejería de Educación JCyL. GIR-SA136G18, GIR-USAL: “Plasticidad, degeneración y regeneración del sistema visual”, Centro en Red de Medicina Regenerativa y Terapia Celular de la JCyL

## Bibliography

Bai, Q., Sun, M., Stolz, D.B., Burton, E.A., 2011. Major isoform of zebrafish P0 is a 23.5 kDa myelin glycoprotein expressed in selected white matter tracts of the central nervous system. J. Comp. Neurol. 519, 1580–1596. https://doi.org/10.1002/cne.22587

Baumann, N., Pham-Dinh, D., 2001. Biology of oligodendrocyte and myelin in the mammalian central nervous system. Physiol. Rev. 81, 871–927.

Bernardos, R.L., Raymond, P.A., 2006. GFAP transgenic zebrafish. Gene Expr. Patterns 6, 1007–1013. https://doi.org/10.1016/j.modgep.2006.04.006

Blasky, A.J., Pan, L., Moens, C.B., Appel, B., 2014. Pard3 regulates contact between neural crest cells and the timing of Schwann cell differentiation but is not essential for neural crest migration or myelination. Dev. Dyn. 243, 1511–23. https://doi.org/10.1002/dvdy.24172

Bribián, A., Barallobre, M.J., Soussi-Yanicostas, N., de Castro, F., 2006. Anosmin-1 modulates the FGF-2-dependent migration of oligodendrocyte precursors in the developing optic nerve. Mol. Cell. Neurosci. 33, 2–14. https://doi.org/S1044-7431(06)00102-3 [pii] 10.1016/j.mcn.2006.05.009

Brösamle, C., 2010. The myelin proteolipid DMalpha in fishes. Neuron Glia Biol. 6, 109–112. https://doi.org/10.1017/S1740925X09000131

Brösamle, C., Halpern, M.E., 2002. Characterization of myelination in the developing zebrafish. Glia 39, 47–57. https://doi.org/10.1002/glia.10088

Buckley, C.E., Goldsmith, P., Franklin, R.J.M., 2008. Zebrafish myelination: a transparent model for remyelination? Dis. Model. Mech. 1, 221–228. https://doi.org/10.1242/dmm.001248

Buckley, C.E., Marguerie, A., Alderton, W.K., Franklin, R.J.M., 2010. Temporal dynamics of myelination in the zebrafish spinal cord. Glia 58, 802–812. https://doi.org/10.1002/glia.20964

Bylund, M., Andersson, E., Novitch, B.G., Muhr, J., 2003. Vertebrate neurogenesis is counteracted by Sox1-3 activity. Nat. Neurosci. 6, 1162–1168. https://doi.org/10.1038/nn1131

Chung, A.-Y., Kim, P.-S., Kim, S., Kim, E., Kim, D., Jeong, I., Kim, H.-K., Ryu, J.-H., Kim, C.-H., Choi, J., Seo, J.-H., Park, H.-C., 2013. Generation of demyelination models by targeted ablation of oligodendrocytes in the zebrafish CNS. Mol. Cells 36, 82–87. https://doi.org/10.1007/s10059-013-0087-9

Czopka, T., 2015. Insights into mechanisms of central nervous system myelination using zebrafish. Glia 64, 333–349. https://doi.org/10.1002/glia.22897

Czopka, T., ffrench-Constant, C., Lyons, D.A., 2013. Individual Oligodendrocytes Have Only a Few Hours in which to Generate New Myelin Sheaths In Vivo. Dev. Cell 25, 599–609. https://doi.org/10.1016/j.devcel.2013.05.013

Czopka, T., Lyons, D.A., 2011. Dissecting Mechanisms of Myelinated Axon Formation Using Zebrafish. Methods Cell Biol. 105, 25–62. https://doi.org/10.1016/B978-0-12-381320-6.00002-3

DeOliveira-Mello L., Lara J.M., Arevalo R., Velasco A., Mack A.F., 2019. Sox2 expression in the visual system of two teleost species. Brain Res, 1722:146350. doi: 10.1016/j.brainres.2019.146350.

Dutton, J.R., Antonellis, A., Carney, T.J., Rodrigues, F.S., Pavan, W.J., Ward, A., Kelsh, R.N., 2008. An evolutionarily conserved intronic region controls the spatiotemporal expression of the transcription factor Sox10. BMC Dev. Biol. 8, 105. https://doi.org/10.1186/1471-213X-8-105

Fang, Y., Lei, X., Li, X., Chen, Y., Xu, F., Feng, X., Wei, S., Li, Y., 2015. A novel model of demyelination and remyelination in a GFP-transgenic zebrafish. Biol. Open 4, 62–68. https://doi.org/10.1242/bio.201410736

Ferilli, M. A. N., Paparella R., Morandini I., Papetti L., Talamanca L. F., Ruscitto C., Ursitti F., Moavero F., Sforza G., Tarantino S., Checchi M. P., Vigevano F., Valeriani M., 2021. Pediatric Neuromyelitis Optica Spectrum Disorder: Case Series and Literature Review. Life (Basel) 12(1):19. doi: 10.3390/life12010019.

Fimbel, S.M., Montgomery, J.E., Burket, C.T., Hyde, D.R., 2007. Regeneration of Inner Retinal Neurons after Intravitreal Injection of Ouabain in Zebrafish. J. Neurosci. 27, 1712–1724. https://doi.org/27/7/1712 [pii] 10.1523/JNEUROSCI.5317-06.2007

Fischer, A.J., Zelinka, C., Scott, M.A., 2010. Heterogeneity of glia in the retina and optic nerve of birds and mammals. PLoS One 5, e10774. https://doi.org/10.1371/journal.pone.0010774

Fitzgibbon, T., Nestorovski, Z., 1997. Morphological Consequences of Myelination in the Human Retina. Exp. Eye Res. 65, 809–819. https://doi.org/10.1006/EXER.1997.0388

Gao, L., Miller, R.H., 2006. Specification of optic nerve oligodendrocyte precursors by retinal ganglion cell axons. J. Neurosci. 26, 7619–7628. https://doi.org/26/29/7619 [pii] 10.1523/JNEUROSCI.0855-06.2006

Garcion, E., Faissner, A., ffrench-Constant, C., 2001. Knockout mice reveal a contribution of the extracellular matrix molecule tenascin-C to neural precursor proliferation and migration. Development 128, 2485–96.

Germanà, A., Montalbano, G., Guerrera, M.C., Amato, V., Laurà, R., Magnoli, D., Campo, S., Suarez-Fernandez, E., Ciriaco, E., Vega, J. a., 2010. Developmental changes in the expression of sox2 in the zebrafish brain. Microsc. Res. Tech. 354, 347–54. https://doi.org/10.1002/jemt.20915

Graham V., Khudyakov J., Ellis P., Pevny L., 2003. SOX2 functions to maintain neural progenitor identity. Neuron, 39(5):749–65. doi: 10.1016/s0896-6273(03)00497-5.

Haldin, C.E., LaBonne, C., 2010. SoxE factors as multifunctional Neural Crest Regulatory Factors. Int. J. Biochem. Cell Biol. 42, 441–444. https://doi.org/10.1038/jid.2014.371

Hines, J.H., Ravanelli, A.M., Schwindt, R., Scott, E.K., Appel, B., 2015. Neuronal activity biases axon selection for myelination in vivo. Nat. Neurosci. 18, 683–689. https://doi.org/10.1038/nn.3992

Hughes, A.N., Appel, B., 2020. Microglia phagocytose myelin sheaths to modifydevelopmental myelination. Nat. Neuroscience. 23, 1055–1066 https://doi.org/10.1038/s41593-020-0654-2

Johns, P.R., Easter Jr., S.S., 1977. Growth of the adult goldfish eye. II. Increase in retinal cell number. J. Comp. Neurol. 176, 331–341. https://doi.org/10.1002/cne.901760303

Jung, S.-H., Kim, S., Chung, A.-Y., Kim, H.-T., So, J.-H., Ryu, J., Park, H.-C., Kim, C.-H., 2010. Visualization of myelination in GFP-transgenic zebrafish. Dev. Dyn. 239, 592–597. https://doi.org/10.1002/dvdy.22166

Kimmel, C.B., Ballard, W.W., Kimmel, S.R., Ullmann, B., Schilling, T.F., 1995. Stages of embryonic development of the zebrafish. Dev. Dyn. 203, 253–310. https://doi.org/10.1002/aja.1002030302

Kordes, U., Cheng, Y.C., Scotting, P.J., 2005. Sox group E gene expression distinguishes different types and maturational stages of glial cells in developing chick and mouse. Brain Res. Dev. Brain Res. 157, 209–213. https://doi.org/S0165-3806(05)00104-5 [pii] 10.1016/j.devbrainres.2005.03.009

Kucenas, S., Snell, H., Appel, B., 2009. nkx2.2a promotes specification and differentiation of a myelinating subset of oligodendrocyte lineage cells in zebrafish. Neuron Glia Biol. 4, 71. https://doi.org/10.1017/S1740925X09990123

Lillo, C., Velasco, A., Jimeno, D., Lara, J.M., Aijón, J., 1998. Ultrastructural organization of the optic nerve of the tench (Cyprinidae, Teleostei). J. Neurocytol. 27, 593–604.

Lyons, D.A., Talbot, W.S., 2015. Glial Cell Development and Function in Zebrafish. Cold Spring Harb. Perspect. Biol. 7, a020586. https://doi.org/10.1101/cshperspect.a020586

Mathews, E.S., Appel, B., 2016. Oligodendrocyte differentiation. Methods Cell Biol. 134, 69–96. https://doi.org/10.1016/bs.mcb.2015.12.004

Mathews, E.S., Mawdsley, D.J., Walker, M., Hines, J.H., Pozzoli, M., Appel, B., 2014. Mutation of 3-Hydroxy-3-Methylglutaryl CoA Synthase I Reveals Requirements for Isoprenoid and Cholesterol Synthesis in Oligodendrocyte Migration Arrest, Axon Wrapping, and Myelin Gene Expression. J. Neurosci. 34, 3402–3412. https://doi.org/10.1523/JNEUROSCI.4587-13.2014

Merchán, P., Bribián, A., Sanchez-Camacho, C., Lezameta, M., Bovolenta, P., de Castro, F., 2007. Sonic hedgehog promotes the migration and proliferation of optic nerve oligodendrocyte precursors. Mol. Cell. Neurosci. 36, 355–368. https://doi.org/S1044-7431(07)00174-1 [pii] 10.1016/j.mcn.2007.07.012

Mercurio S., Serra L., Motta A., Gesuita L, Sanchez-Arrones L., Inverardi F., Foglio B., Barone C., Kaimakis P., Martynoga B., Ottolenghi S., Studer M., Guillemot F., Frassoni C., Bovolenta P., Nicolis S.K., 2019. Sox2 Acts in Thalamic Neurons to Control the Development of Retina-Thalamus-Cortex Connectivity. iScience, 15:257–273. doi: 10.1016/j.isci.2019.04.030.

Miller, R.H., 2002. Regulation of oligodendrocyte development in the vertebrate CNS. Prog. Neurobiol. 67, 451–67.

Münzel, E.J., Becker, C.G., Becker, T., Williams, A., 2014. Zebrafish regenerate full thickness optic nerve myelin after demyelination, but this fails with increasing age. Acta Neuropathol. Commun. 2, 77. https://doi.org/10.1186/s40478-014-0077-y

Münzel, E.J., Schaefer, K., Obirei, B., Kremmer, E., Burton, E. a., Kuscha, V., Becker, C.G., Brösamle, C., Williams, A., Becker, T., 2012. Claudin k is specifically expressed in cells that form myelin during development of the nervous system and regeneration of the optic nerve in adult zebrafish. Glia 60, 253–270. https://doi.org/10.1002/glia.21260

Nakamura, K., Harada, C., Namekata, K., Harada, T., 2006. Expression of olig2 in retinal progenitor cells. Neuroreport 17, 345–349. https://doi.org/10.1097/01.wnr.0000203352.44998.6b

Nawaz, S., Schweitzer, J., Jahn, O., Werner, H.B., 2013. Molecular evolution of myelin basic protein, an abundant structural myelin component. Glia 61, 1364–1377. https://doi.org/10.1002/glia.22520

Ono, K., Yasui, Y., Rutishauser, U., Miller, R.H., 1997. Focal ventricular origin and migration of oligodendrocyte precursors into the chick optic nerve. Neuron 19, 283–292. https://doi.org/S0896-6273(00)80939-3 [pii]

Ono, K., Yoshii, K., Tominaga, H., Gotoh, H., Nomura, T., Takebayashi, H., Ikenaka, K., 2017. Oligodendrocyte precursor cells in the mouse optic nerve originate in the preoptic area. Brain Struct. Funct. https://doi.org/10.1007/s00429-017-1394-2

Park, H.-C., Mehta, A., Richardson, J.S., Appel, B., 2002. olig2 is required for zebrafish primary motor neuron and oligodendrocyte development. Dev. Biol. 248, 356–368. https://doi.org/10.1006/dbio.2002.0738

Parrilla, M., León-Lobera, F., Lillo, C., Arévalo, R., Aijón, J., Lara, J.M., Velasco, A., 2016. Sox10 Expression in Goldfish Retina and Optic Nerve Head in Controls and after the Application of Two Different Lesion Paradigms. PLoS One 11, e0154703. https://doi.org/10.1371/journal.pone.0154703

Parrilla, M., Lillo, C., Herrero-Turrion, M.J., Arevalo, R., Aijon, J., Lara, J.M., Velasco, A., 2013. Pax2+ astrocytes in the fish optic nerve head after optic nerve crush. Brain Res. 1492, 18–32. https://doi.org/10.1016/j.brainres.2012.11.014

Parrilla, M., Lillo, C., Herrero-Turrion, M.J., Arevalo, R., Aijon, J., Lara, J.M., Velasco, A., 2012. Characterization of Pax2 expression in the goldfish optic nerve head during retina regeneration. PLoS One 7, e32348. https://doi.org/10.1371/journal.pone.0032348

Pinzon-Olejua, A., Welte, C., Chekuru, A., Bosak, V., Brand, M., Hans, S., Stuermer, C.A.O., 2017. Cre-inducible site-specific recombination in zebrafish oligodendrocytes. Dev. Dyn. 246, 41–49. https://doi.org/10.1002/dvdy.24458

Plavicki, J.S., Baker, T.R., Burns, F.R., Xiong, K.M., Gooding, A.J., Hofsteen, P., Peterson, R.E., Heideman, W., 2014. Construction and characterization of a sox9b transgenic reporter line. Int. J. Dev. Biol. 58, 693–699. https://doi.org/10.1387/ijdb.140288jp

Preston, M.A., Macklin, W.B., 2015. Zebrafish as a model to investigate CNS myelination. Glia 63, 177–193. https://doi.org/10.1002/glia.22755

Ravanelli, A.M., Appel, B., 2015. Motor neurons and oligodendrocytes arise from distinct cell lineages by progenitor recruitment. Genes Dev. 29, 1–12. https://doi.org/10.1101/gad.271312.115

Ravanelli, A.M., Kearns, C.A., Powers, R.K., Wang, Y., Hines, J.H., Donaldson, M.J., Appel, B., 2018. Sequential specification of oligodendrocyte lineage cells by distinct levels of Hedgehog and Notch signaling. Dev. Biol. https://doi.org/10.1016/j.ydbio.2018.10.004

Reichenbach, A., Bringmann, A., 2015. Retinal glia, Encyclopedia of Neuroscience. Morgan & Claypool Life Sciences. https://doi.org/10.4199/C00122ED1V01Y201412NGL003

Reichenbach, A., Schippel, K., Schumann, R., Hagen, E., 1988. Ultrastructure of rabbit retinal nerve fibre layer--neuro-glial relationships, myelination, and nerve fibre spectrum. J. Hirnforsch. 29, 481–491.

Reinhardt R., Centanin L., Tavhelidse T., Inoue D., Wittbrodt B., Concordet JP., Martinez-Morales J.R., Wittbrodt J., 2015. Sox2, Tlx, Gli3, and Her9 converge on Rx2 to define retinal stem cells in vivo. EMBO J, 34(11):1572–88. doi: 10.15252/embj.201490706.

Santos-Ledo, A., Garcia-Macia, M., Campbell, P.D., Gronska, M., Marlow, F.L., 2017. Kinesin-1 promotes chondrocyte maintenance during skeletal morphogenesis. PLoS Genet 13(7): e1006918. https://doi.org/10.1371/journal.pgen.1006918

Schweitzer, J., Becker, T., Becker, C.G., Schachner, M., 2003. Expression of protein zero is increased in lesioned axon pathways in the central nervous system of adult zebrafish. Glia 41, 301–317. https://doi.org/10.1002/glia.10192

Shibasaki, K., Takebayashi, H., Ikenaka, K., Feng, L., Gan, L., 2007. Expression of the basic helix-loop-factor Olig2 in the developing retina: Olig2 as a new marker for retinal progenitors and late-born cells. Gene Expr. Patterns 7, 57–65. https://doi.org/10.1016/j.modgep.2006.05.008

Shin, J., Park, H.-C., Topczewska, J.M., Mawdsley, D.J., Appel, B., 2003. Neural cell fate analysis in zebrafish using olig2 BAC transgenics. Methods Cell Sci. 25, 7–14. https://doi.org/10.1023/B:MICS.0000006847.09037.3a

Stolt, C.C., Wegner, M., 2010. SoxE function in vertebrate nervous system development. Int. J. Biochem. Cell Biol. 42, 437–440. https://doi.org/10.1016/j.biocel.2009.07.014

Takada, N., Appel, B., 2010. Identification of genes expressed by zebrafish oligodendrocytes using a differential microarray screen. Dev. Dyn. 239, 2041–2047. https://doi.org/10.1002/dvdy.22338

Takada, N., Kucenas, S., Appel, B., 2010. Sox10 is necessary for oligodendrocyte survival following axon wrapping. Glia 58, 996–1006. https://doi.org/10.1002/glia.20981

Thummel, R., Kassen, S.C., Enright, J.M., Nelson, C.M., Montgomery, J.E., Hyde, D.R., 2008. Characterization of Müller glia and neuronal progenitors during adult zebrafish retinal regeneration. Exp. Eye Res. 87, 433–444. https://doi.org/10.1016/j.exer.2008.07.009

Tian, C., Zou, S., Hu, B., A., W., RA., N., PA., R., 2016. Extraocular Source of Oligodendrocytes Contribute to Retinal Myelination and Optokinetic Responses in Zebrafish. Investig. Opthalmology Vis. Sci. 57, 2129. https://doi.org/10.1167/iovs.15-17675

Tiwari, S., Dharmarajan, S., Shivanna, M., Otteson, D.C., Belecky-Adams, T.L., 2014. Histone deacetylase expression patterns in developing murine optic nerve. BMC Dev. Biol. 14, 30. https://doi.org/10.1186/1471-213X-14-30

Trost, A., Schroedl, F., Lange, S., Rivera, F.J., Tempfer, H., Korntner, S., Stolt, C.C., Wegner, M., Bogner, B., Kaser-Eichberger, A., Krefft, K., Runge, C., Aigner, L., Reitsamer, H.A., 2013. Neural crest origin of retinal and choroidal pericytes. Investig. Ophthalmol. Vis. Sci. 54, 7910–7921. https://doi.org/10.1167/iovs.13-12946

Yokoi, H., Yan, Y.-L., Miller, M.R., BreMiller, R.A., Catchen, J.M., Johnson, E.A., Postlethwait, J.H., 2009. Expression profiling of zebrafish sox9 mutants reveals that Sox9 is required for retinal differentiation. Dev. Biol. 329, 1–15. https://doi.org/10.1016/j.ydbio.2009.01.002

Yoshida, M., Macklin, W.B., 2005. Oligodendrocyte development and myelination in GFP-transgenic zebrafish. J. Neurosci. Res. 81, 1–8. https://doi.org/10.1002/jnr.20516

